# Persistent CAD activity in memory CD8+ T cells supports rRNA synthesis and ribosomal biogenesis required at rechallenge

**DOI:** 10.1101/2021.11.10.468070

**Authors:** Michael D. Claiborne, Srona Sengupta, Liang Zhao, Matthew L. Arwood, Im-Meng Sun, Jiayu Wen, Elizabeth A. Thompson, Marisa Mitchell-Flack, Marikki Laiho, Jonathan D. Powell

## Abstract

Memory CD8+ T cells are characterized by their ability to persist long after the initial antigen encounter and their ability to generate a rapid recall response. Recent studies have identified a role for metabolic reprogramming and mitochondrial function in promoting the longevity of memory T cells. However, detailed mechanisms involved in promoting the rapid recall response are incompletely understood. Here we identify a novel role for the initial and continued activation of the trifunctional rate-limiting enzyme of the *de novo* pyrimidine synthesis pathway CAD (carbamoyl-phosphate synthetase 2, aspartate transcarbamylase, and dihydroorotase) as critical in promoting the rapid recall response of previously-activated CD8+ T cells. CAD is rapidly phosphorylated upon T cell activation in an mTORC1-dependent manner yet remains phosphorylated long after initial activation. Previously-activated CD8+ T cells display continued *de novo* pyrimidine synthesis in the absence of mitogenic signals and interfering with this pathway diminishes the speed and magnitude of cytokine production upon rechallenge. Inhibition of CAD does not affect cytokine transcript levels, but diminishes available pre-rRNA, the polycistronic rRNA precursor whose synthesis is the rate-limiting step in ribosomal biogenesis. CAD inhibition additionally decreases levels of detectable ribosomal proteins in previously-activated CD8+ T cells. Overexpression of CAD improves both the cytokine response and proliferation of memory T cells. Overall, our studies reveal a novel and critical role for CAD-induced pyrimidine synthesis and ribosomal biogenesis in promoting the rapid recall response characteristic of memory T cells.

**One Sentence Summary:** Pyrimidine synthesis fuels ribosomal biogenesis to facilitate rapid recall responses in CD8+ T cells

## Introduction

Memory CD8+ T cells are characterized by two important functional properties. They persist at relatively increased frequencies long after their initial antigen encounter and, upon rechallenge, memory T cells exhibit both a rapid and robust recall response (*1,2*). While previous studies have identified metabolic reprogramming as promoting persistence (*3*), less is known about the biochemical properties that endow memory CD8+ cells with superior recall capacity. The kinase mTOR, acting through its two canonical signaling complexes mTORC1 and mTORC2, integrates environmental cues to direct a myriad of transcriptional and translational programs downstream of T cell activation (*4,5*). Specifically, mTORC1 promotes CD8+ effector generation and mTORC2 inhibits the generation of memory CD8+ T cells, although the downstream targets of mTOR giving rise to these different T cell fates have not been fully elucidated (*6*). To this end, asymmetric partitioning of mTOR activity promotes in part the simultaneous generation of effector and memory T cells during an immune response (*7*).

Naïve T cells maintain low baseline levels of mTOR activity and divide infrequently without TCR engagement during homeostatic proliferation. Upon T cell activation, the PI3K/Akt/mTOR signaling cascade results in robust mTORC1 activity that is detectable within minutes and returns to baseline by 48-72 hours (*9*). This mTORC1 activity, in concert with other signaling pathways, dramatically alters the physiology and metabolism of lymphocytes to promote a glycolytic metabolism that is permissive for the additional synthesis of necessary metabolites (*5,6*). Recently, mTORC1 acting via the downstream kinase S6K1 was shown to phosphorylate and activate the rate-limiting trifunctional enzyme of the *de novo* pyrimidine synthesis pathway, CAD (carbamoyl-phosphate synthetase 2, aspartate transcarbamylase, and dihydroorotase) in HEK-293 and HeLa cell lines (*10*). In the absence of mitogenic signals, T cells do not require *de novo* pyrimidine synthesis for cytidine and thymidine nucleosides, instead relying on the pyrimidine salvage pathway (*14,15*). The rapid clonal expansion of activated T cells is supported in part by rapid *de novo* synthesis of nucleic acids, consistent with multiple rounds of cell division each requiring replication of genomic DNA. The same radiolabeling studies that helped describe this process over fifty years ago also noted the continued accumulation of tritiated nucleotides into RNA in previously-activated lymphocytes nine days after stimulation -- well after incorporation of new nucleotides into DNA had diminished (*16*), although an explanation for this phenomenon was not explored at the time.

Inasmuch as full T cell activation leading to effector cell generation results in mTORC1 activation and CAD-induced pyrimidine synthesis is mTORC1 dependent, we hypothesized that the ability of mTORC1 to promote effector cell generation was due in part to the acute activation of CAD. Surprisingly, while we did indeed find that TCR engagement leads to CAD phosphorylation, we also observed that phosphorylated CAD remains detectable in resting cells long after T cell activation at a time when mTOR activity had returned to baseline. This prompted us to explore the role of persistent CAD activation in previously activated/memory T cells. Our studies demonstrate that persistent CAD-induced pyrimidine synthesis contributes to the rapid recall response of memory T cells. Mechanistically, the contribution of CAD is not mediated through increased cytokine transcription, but rather promotion of ribosomal biogenesis secondary to CAD-mediated pre-rRNA synthesis.

## Results

### CAD undergoes TORC1-mediated phosphorylation upon CD8+ T cell activation and remains phosphorylated in resting cells

The trifunctional CAD protein catalyzes the first three steps in *de novo* pyrimidine synthesis (Figure 1A). Regulation of CAD activity by phosphorylation has been described at multiple residues by a diverse array of kinases (*8*), including an activating phosphorylation at Ser1859 by mTORC1 acting via the downstream kinase S6K1 (*10*). As mTOR signaling is instrumental in CD8+ T cell activation and differentiation (*5,6*), we interrogated CD8+ T cells for CAD phosphorylation via Western blot at this site following T cell activation. CAD is phosphorylated at Ser1859 in a mTORC1-dependent manner acutely following T cell activation, as T cells activated in the presence of the mTORC1 inhibitor rapamycin showed no increase in pCAD or mTOR activity as measured by pS6K1 (Figure 1B). This phosphorylation was further confirmed to be mTORC1-dependent by utilizing CD8+ cells isolated from T cell-specific *Rptor* or *Rictor* knockout mice with T cells deficient in mTORC1 or mTORC2 signaling, respectively (Figure 1C). In these experiments, the elimination of mTORC1 signaling abrogated CAD phosphorylation while genetic inhibition of mTORC2 had no effect. After validating the use of our commercially available pCAD antibody for flow cytometry applications via shRNA knockdown of CAD (Figure S1), we confirmed mTORC1-dependent phosphorylation of CAD upon CD8+ T cell activation utilizing flow cytometry (Figure 1D). Thus, TCR engagement leads to CAD phosphorylation in an mTORC1 dependent fashion.

**Figure 1:**
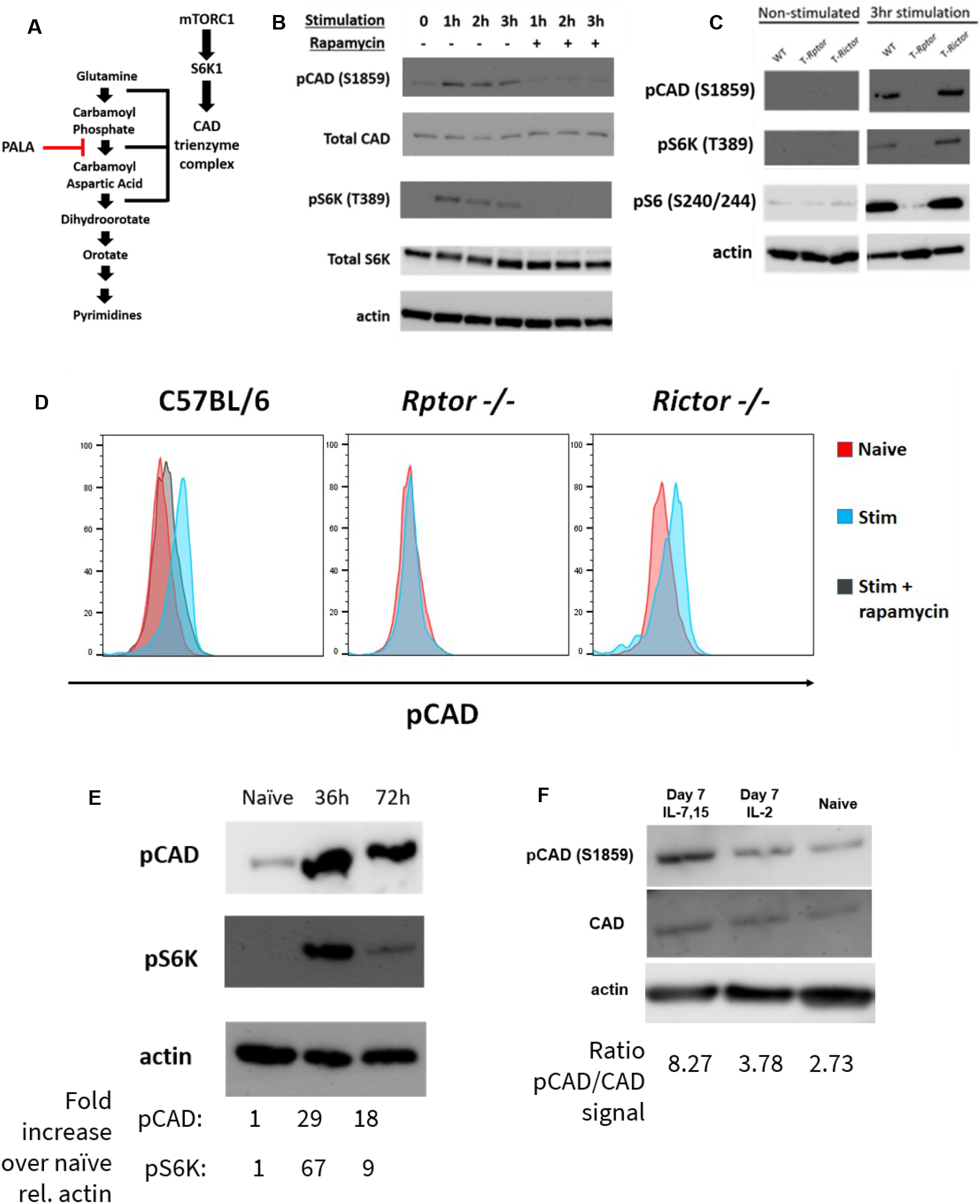
mTORC1-dependent phosphorylation of CAD at S1859 persists beyond T cell activation. (A) The *de novo* pyrimidine synthesis pathway. The rate-limiting trifunctional CAD protein catalyzes the first three steps of this pathway in mammalian cells. (B) Western blot analysis of CD8+ T cells isolated from spleens of naïve C57BL/6 mice and stimulated with α-CD3/28 antibodies for indicated time periods with or without rapamycin (100nM). (C) Western blot analysis of CD8+ T cells isolated from spleens of naïve C57BL/6, *Rptor* and *Rictor* mice stimulated with α-CD3/28 antibodies for indicated time periods. *Rptor*- and *Rictor-floxed* mice were crossed with CD4-Cre mice and backcrossed onto the C57BL/6 background for nine generations. (D) Flow cytometric analysis CD8+ T cells isolated from spleens of naïve C57BL/6, *Rptor* and *Rictor* mice and stimulated with α-CD3/28 antibodies for indicated time periods with or without rapamycin (100nM) before fixation and interrogation for pCAD (S1859). (E) Western blot analysis of CD8+ T cells isolated from spleens of naïve C57BL/6 mice and stimulated with α-CD3/28 antibodies for 36 and 72 hours. (F) Western blot analysis of CD8+ T cells isolated from spleens of naïve C57BL/6 mice and stimulated with α-CD3/28 antibodies for 48 hours, then expanded in media containing IL-2 or IL-7 + IL-15 for an additional five days. Data in B-E are representative of three independent experiments.

Next, we cultured CD8+ T cells for up to 72 hours after activation to further interrogate CAD phosphorylation. At 36 hours after activation, both pCAD and pS6K are robustly elevated. However, by 72 hours of stimulation, mTORC1 activity begins to decrease while pCAD signal remains elevated when compared to naïve cells (Figure 1E). That is, we observed prolonged phosphorylation of CAD in previously-activated T cells long after mTORC1 activity diminished. This continuation of pCAD signal led us to hypothesize that elevated pCAD might not just be important for acute activation of T cells, but also may play a role in the identity and function of long-lived memory CD8+ T cells. Expansion of activated CD8+ T cells with the cytokines IL-7 and IL-15 induces a memory-like phenotype *in vitro* when compared to expansion with IL-2, which induces an effector-like phenotype (*11*). Indeed, CD8+ T cells cultured for seven days following activation have elevated pCAD compared to naïve cells, with activated cells cultured in IL-7 and IL-15 being further enriched in pCAD than cells cultured in IL-2 (Figure 1F). Taken together, these findings identify Ser1859 of CAD as a downstream target of mTORC1 in CD8+ T cells. Upon T cell activation, this site is phosphorylated in an mTORC1-dependent manner yet remains phosphorylated *in vitro* even after mTORC1 activity decreases.

### CD8+ T cells maintain an activation-induced increase in *de novo* pyrimidine synthesis without further stimulation

To confirm whether the elevated pCAD seen in previously-activated CD8+ T cells reflected an increase in *de novo* pyrimidine synthesis, we utilized a mass spectrometry approach to quantify metabolites in this pathway. Consistent with previous findings, naïve T cells generated limited amounts of carbamoyl aspartic acid, orotate, and uridine monophosphate (UMP), three intermediates of *de novo* pyrimidine synthesis (Figure 2A). Activation of naïve T cells for 24 hours led to an increase in these metabolites consistent with prior studies (17). To further interrogate the fates of metabolic intermediates in the *de novo* pyrimidine synthesis pathway, we quantified metabolic flux utilizing amide-labeled ^15^-N glutamine (*10*). The protocol is diagrammed in Figure 2B. Of note, as the amide nitrogen of one molecule of glutamine is condensed with CO2 in the first step of this pathway, all downstream metabolites except for CTP will be reflected in the M+1 isotopic peak. As demonstrated in Figure 2C, naïve CD8+ T cells exhibited limited flux into *de novo* pyrimidine synthesis which increased upon TCR stimulation. Surprisingly, previously-activated cells maintained a high baseline level of flux into this pathway even in the absence of further stimulation. These findings demonstrate that the increased pCAD seen in previously-activated CD8+ T cells *in vitro* indeed correlates with an increase in both detectable metabolites in the *de novo* pyrimidine synthesis pathway and flux into this pathway via amide-labeled nitrogen tracing.

**Figure 2:**
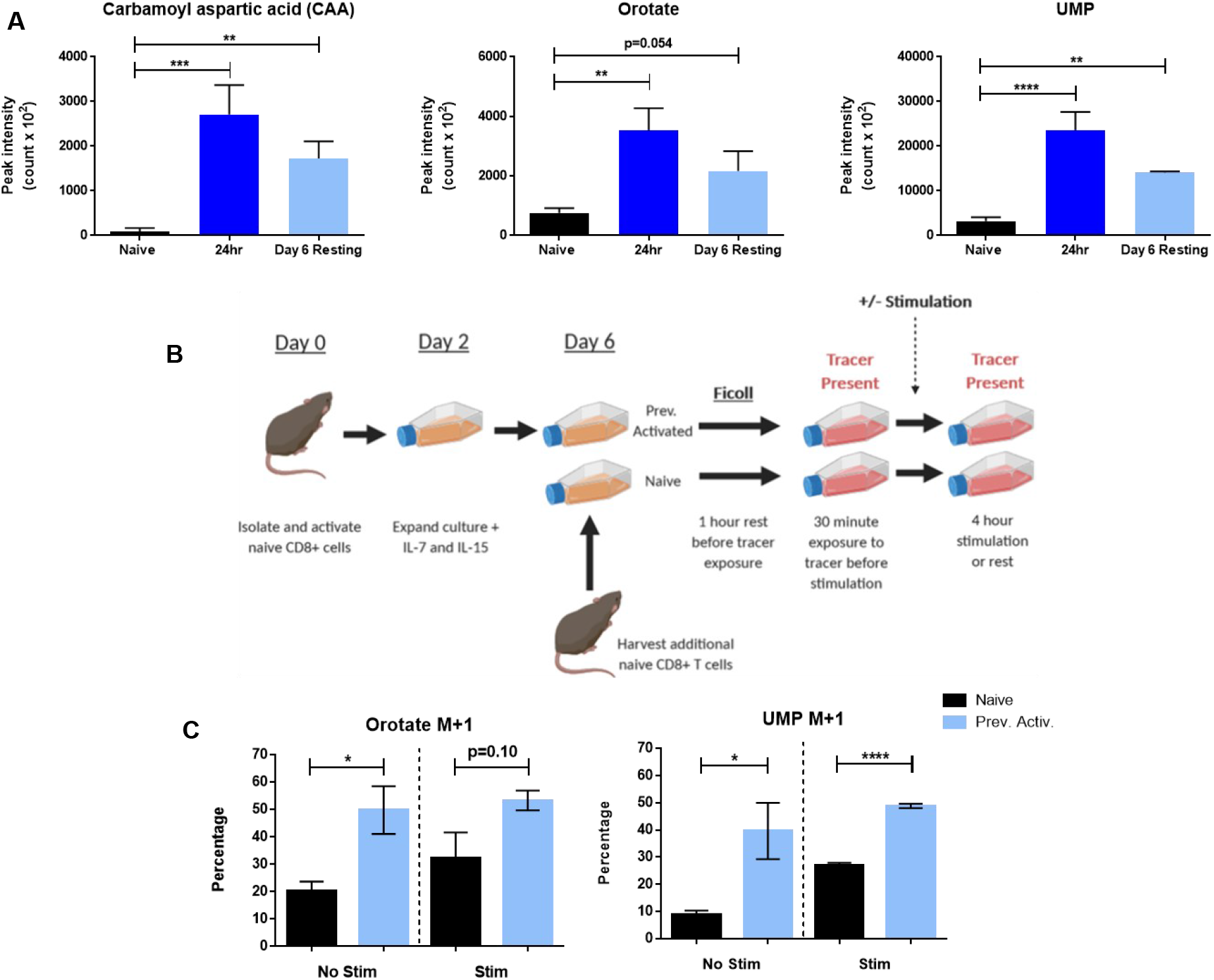
CD8+ T cells continue to conduct de novo pyrimidine synthesis in vitro beyond T cell activation. (A) Peak area for *de novo* pyrimidine synthesis metabolites measured by LC/MS. CD8+ T cells were isolated from spleens of naïve C57BL/6 mice and stimulated with α-CD3/28 antibodies. Cells were expanded at 48 hours into culture with IL-7 + IL-15 and rested four more days to generate Day 6 Resting condition. Additional naïve CD8+ T cells were isolated before analysis (Naïve condition). (B) Schematic of amide-labeled ^15^-N glutamine tracing. Previously Activated cells were generated by stimulating isolated CD8+ cells from naïve C57BL/6 mice with α-CD3/28 antibodies for 48 hours before expanding media with IL-7 + IL-15 and rested until Day 6 of culture, when an additional naïve C57BL/6 mouse was harvested and CD8+ splenocytes isolated. Naïve and Previously Activated cells were then rested before transfer to glutamine-free media supplemented with 4mM amide-labeled ^15^-N glutamine, and a portion of each condition was stimulated with α-CD3/28 antibodies. (C) Percentage M+1 peak data for *de novo* pyrimidine synthesis metabolites measured by LC/MS. SRM values are listed in Supplemental Materials S7. Data in A and C are representative of triplicate samples analyzed from two independent experiments. Data in A show mean ± SD analyzed by one-way ANOVA followed by Sidak’s Multiple Comparison Test. Data in C show mean ± SD analyzed by unpaired Student’s t-test. *p <0.05, **p<0.01, ***p<0.001, ****p<0.0001.

### Phosphorylated CAD is enriched in murine central memory CD8+ cells only with intact mTORC1 signaling at activation

To investigate the phosphorylation status of CAD *in vivo*, we first isolated spleens from nine-week-old uninfected C57BL/6 mice for analysis by flow cytometry (Figure 3A). In the absence of any experimental infection, pCAD was enriched in central memory-phenotype CD8+ cells (CD44^+^ CD62L^+^) over the naïve CD8+ cells in the same animals (CD44^-^CD62L^+^). To quantify pCAD levels in an antigen-specific setting, we isolated CD8+ cells from transgenic OT-I mice overexpressing the TCR specific for OVA_257-264_ (SIINFEKL) epitope. These cells also expressed the congenic marker CD90.1 (Thy1.1), allowing us to adoptively transfer them into CD90.2 (Thy1.2) recipients and infect them with Vaccinia virus expressing OVA (Vac-OVA) to expand a population of antigen-specific cells. Antigen-specific central memory CD8+ cells (CD44^+^CD62L^+^Thy1.1^+^) at Day 45 post-infection were enriched in pCAD relative to naïve CD8+ T cells in each mouse (CD44^-^CD62L^+^Thy1.1^-^) (Figure 3B). To investigate enrichment of pCAD in relation to mTOR activity, we repeated this experiment but harvested splenocytes both at the peak of Vac-OVA infection (Day 8) and during the contraction phase (Day 15). Both mTOR activity (as measured by pS6) and pCAD were elevated in antigen-responding cells compared to endogenous naïve cells (Figure 3C). During the contraction phase, however, pS6 returned to the baseline level seen in naïve cells while pCAD remained enriched in OVA-specific T cells. These findings demonstrate that the persistence of pCAD in the absence of continued mTORC1 activity as seen in previously-activated CD8+ T cells *in vitro* extends to central memory CD8+ T cells *in vivo*.

**Figure 3:**
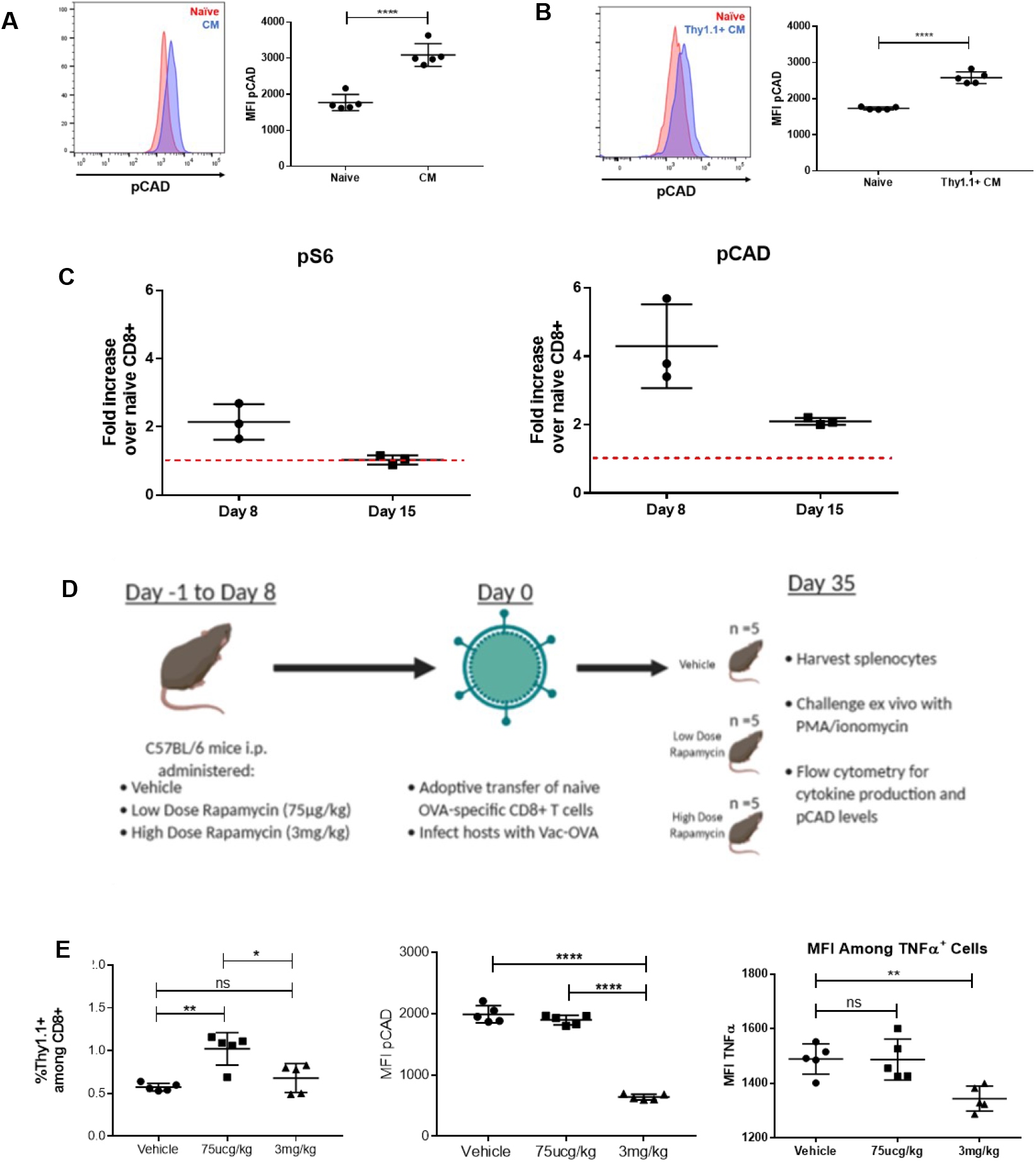
pCAD is detectable in central memory CD8+ cells in vivo when mTORC1 is intact at T cell activation. (A) Flow cytometric analysis of pCAD signal in naïve cells (Live CD8+CD44-CD62L+) and CM cells (Live CD8+CD44+CD62L+) isolated from the spleens of uninfected C57BL/6 mice. (B) Flow cytometric analysis of pCAD in naïve and antigen-specific central memory CD8+ cells. CD8+ cells were isolated from spleens of naïve OT-I mice and adoptively transferred into C57BL/6 hosts at 10^5^ cells/mouse. Hosts were immediately infected with Vac-OVA and splenocytes were harvested forty-five days later to compare pCAD levels in naïve (Live CD8+Thy1.1-) and central memory (Live CD8+Thy1.1+CD62L+) cells. (C) Flow cytometric analysis of pCAD and pS6 during viral infection. Experiment was performed as in 3B with splenocytes harvested eight days and fifteen days post-infection. (D) Schematic of in vivo rapamycin administration. Mice were administered Vehicle, Low Dose (75μg/kg) or High Dose (3mg/kg) rapamycin for Day −1 to Day 8 of Listeria-OVA infection with adoptive transfer of 10^5^ Thy1.1^+^ CD8+ cells freshly isolated from naïve OT-I mice on Day 0. Splenocytes were harvested on Day 35 of infection with a portion restimulated with PMA/ionomycin for 4 hours before analysis by flow cytometry. (E) Splenocytes from experiments described in Figure 3D. A, B, and E are representative of three independent experiments of n=5 mice per experiment. C is representative of three independent experiments with n = 3 mice per condition. Data in A and B show mean fluorescence intensity values ± SD analyzed by unpaired Student’s t-test. Data in E show percentage of Thy1.1+ cells among CD8+ cells or mean fluorescence intensity values ± SD analyzed by one-way ANOVA followed by Sidak’s Multiple Comparison Test. *p <0.05, **p<0.01, ***p<0.001, ****p<0.0001, ns = not significant.

Our data thus far demonstrate that upon initial activation, TCR-induced mTORC1 activation leads to phosphorylation of CAD, which in turn promotes memory recall responses. At first glance, these data contradict previous studies demonstrating increased memory T cell generation in the presence of rapamycin (*12*). To further clarify the role of mTORC1 signaling in CAD phosphorylation, we modified the experimental design utilized by Araki et al. to evaluate the effect of mTOR inhibition during the naïve-to-effector transition of CD8+ cells responding to infection (*12*). We adoptively transferred OVA-specific Thy1.1^+^ CD8+ T cells into wild-type recipients and infected them with Vaccinia-OVA (Figure 3D). The mice were administered low-dose rapamycin (75μg/kg), high-dose rapamycin (3mg/kg), or vehicle from day −1 to day 8 of infection and the spleens were harvested on day 35. A portion of splenocytes from each mouse was fixed immediately for analysis of pCAD levels whereas another portion was restimulated for 4 hours before fixation to interrogate for cytokine production. Consistent with previous findings (*12*), low-dose rapamycin administered during the naïve-to-effector transition increased the frequency of memory cells. However, this low dose does not inhibit the phosphorylation of CAD when compared to vehicle controls (Figure 3E). Thus, our findings concerning the role of pCAD in promoting memory T cell function are compatible with the previously established model in which low-dose rapamycin promotes memory T cell formation (*13*). Alternatively, administration of high-dose rapamycin during this time decreased detectable pCAD. When these memory cells were restimulated, rapamycin treatment increased the percentage TNFα and IL-2-positive cells, with low-dose treatment resulting in a greater increase than high-dose treatment (Figure S2). However, with increasing doses of rapamycin, the MFI of cytokine signal among cytokine-positive cells decreased (Figure 3E, S2). Taken with Figure 1, these data suggest pCAD is enriched in previously-activated cells *in vivo* and *in vitro* and this enrichment is abrogated in situations where mTORC1 is inhibited at T cell activation. pCAD levels seen in antigen-specific memory cells are associated with the ability of those cells to produce cytokine upon rechallenge *ex vivo*.

### Reduction of CAD activity before rechallenge decreases the speed and magnitude of CD8+ recall responses

Blasting CD8+ cells are known to require nucleotides produced by *de novo* pyrimidine synthesis (*15,17*). Thus, we hypothesized that the CAD-induced increase in *de novo* pyrimidine synthesis promoted the rapid recall response. To this end, we sought to inhibit CAD activity during the resting phase prior to rechallenge. We activated and expanded antigen-specific CD8+ cells and treated them with the CAD inhibitor N-phosphonacetyl-L-aspartate (PALA), a highly selective competitive inhibitor of CAD (*18*, Figure 1A). To confirm PALA’s activity as a CAD inhibitor, we treated previously-activated cells with PALA for the final 24 hours of their six day culture period, washed out any remaining drug, then exposed them to amide-labeled ^15^-N glutamine-containing media alongside naïve cells with and without TCR stimulation. PALA-pretreatment decreased flux into *de novo* pyrimidine synthesis in both resting cells and rechallenged cells in the absence of continued drug exposure (Figure 4A). PALA did not cause global metabolic inhibition, as PALA treatment moderately increased flux into unrelated pathways such as the *de novo* purine synthesis pathway in cells at rest (Figure S3). These results confirm PALA as a specific inhibitor of *de novo* pyrimidine synthesis for use in our studies.

**Figure 4:**
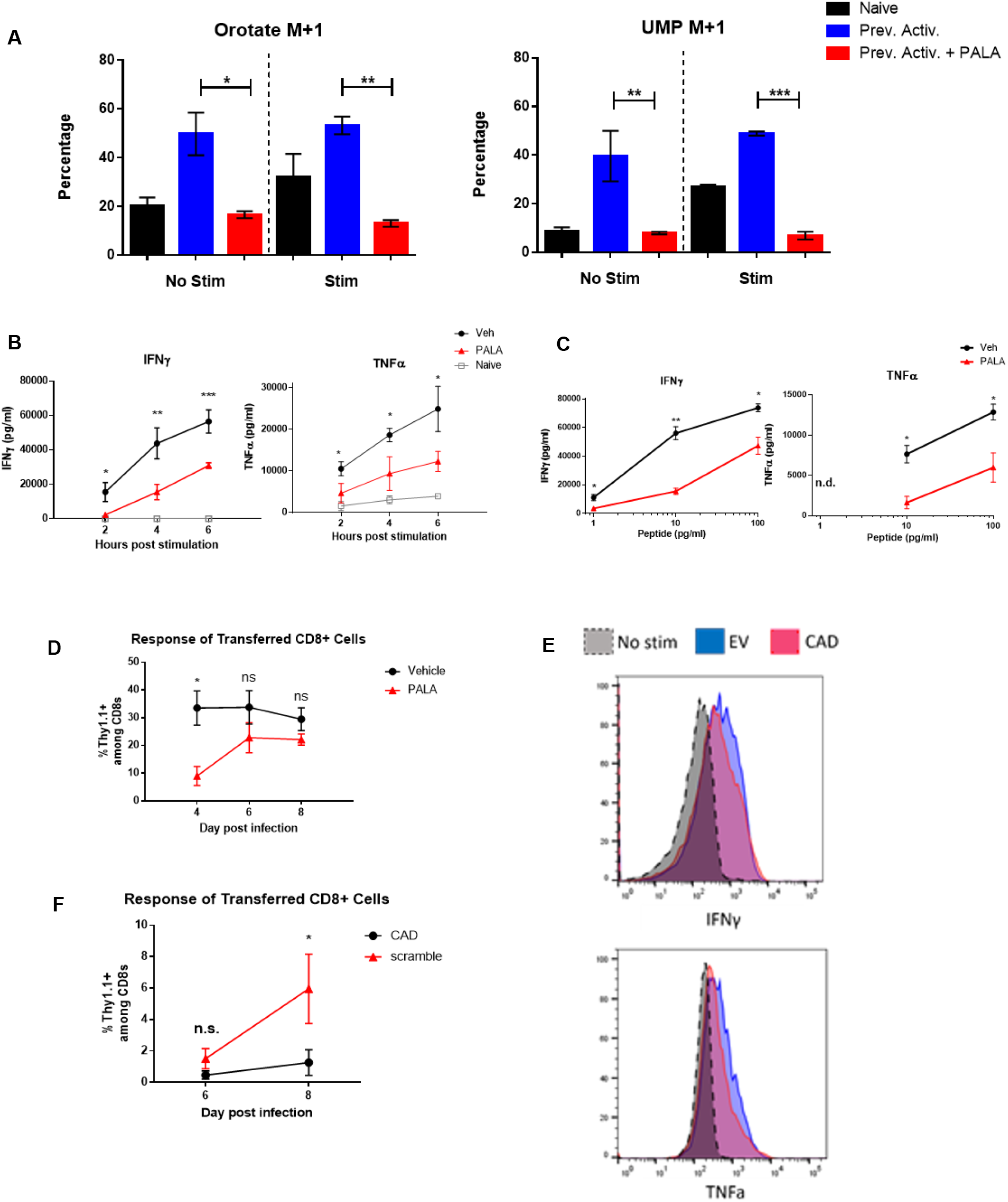
CAD inhibition in resting cells impairs their response at rechallenge. (A) Percentage M+1 peak data for *de novo* pyrimidine synthesis metabolites measured by LC/MS. Previously Activated cells were generated as in Figure 2C, with the addition of Vehicle or 1mM PALA for the final 24 hours of culture. Naïve and Previously Activated cells were rested in culture media before transfer to glutamine-free media supplemented with 4mM amide-labeled ^15^-N glutamine, and a portion of each condition was stimulated with α-CD3/28 antibodies. Naïve and Previously Activated cells analyzed are the same triplicate samples from two independent experiments utilized in Figure 2C. (B) Cytokine response of previously activated cells over a time course of restimulation as measured by ELISA. Vehicle and PALA-treated cells were generated as in Figure 4A. On Day 6 of the experiment, naïve OT-I splenocytes were harvested, irradiated, and incubated with OVA_257-264_ (SIINFEKL) peptide for 45 minutes before plating with cultured T cells. (C) Cytokine response of previously activated cells to decreasing peptide doses as measured by ELISA. Vehicle and PALA-treated cells were generated as in Figure 4A from OT-I mice. On Day 6 of experiment, naïve OT-I splenocytes were harvested, irradiated, and incubated with indicated dose of OVA_257-264_ (SIINFEKL) peptide for 45 minutes before plating with cultured T cells. Supernatants harvested and analyzed for cytokine by ELISA at 5 hours, n.d. equals not detected. (D) Response of transferred Previously Activated p14 cells to LCMV-Armstrong infection with and without PALA pretreatment. Mice were cheek bled on indicated days with percentage of Thy1.1 ^+^ cells among CD8+ cells in blood quantified by flow cytometry. (E) Flow cytometric analysis of cytokine production of Previously Activated cells following lentiviral transduction with *Cad*-targeted or scramble viral constructs. Non-stimulated cells shown in grey. (F). Response of transferred Previously Activated P14 cells to LCMV-Armstrong infection with and without shRNA knockdown of CAD. Mice were cheek bled on indicated days in an analogous manner to Figure 3D. Data in A representative of two independent experiments. Data in B and C are mean concentration ± SD of two to three dilutions of supernatant each from three independent experiments. Data from D representative of three independent experiments with n = 5 mice per condition. Data from E representative of two independent experiments. Data in B, C, D, and F are analyzed by unpaired Student’s t-test at each indicated time point or peptide concentration. *p <0.05, **p<0.01, ***p<0.001, ns = not significant.

We next sought to determine the functional consequences of CAD inhibition. Previously-activated cells were generated as described, and PALA was added for 24 hours on the sixth day of culture. Next the PALA was washed out and dead cells were removed by Ficoll gradient prior to rechallenge utilizing irradiated murine antigen presenting cells pre-loaded with peptide. As expected, previously-activated cells produced more TNFα and IFNγ faster than naïve cells, with IFNγ undetectable in newly activated naïve cells (Figure 4B). However, PALA-pretreatment decreased the speed with which previously-activated cells were able to produce both TNFα and IFNγ. Of note is the fact that PALA is not present during rechallenge, and thus these observations reflect inhibition of CAD activity during the baseline rest period.

In addition to a more robust and rapid response to antigen, memory T cells can also respond robustly to lower doses of antigen (*19*). Thus, we hypothesized that inhibition of CAD in memory cells would lead to a decrease in the dose response curve of these cells. To address this, we generated previously-activated cells pretreated with PALA as in Figure 4A and exposed them to irradiated antigen-presenting cells loaded with peptide, with supernatants harvested at 5 hours. PALA pretreatment decreased the ability of previously-activated cells to respond to low concentrations of antigen (Figure 4C). Overall, our results suggest that inhibition of CAD following initial T cell expansion -but prior to rechallenge - impairs both the speed and magnitude of the CD8+ recall response.

To explore the effects of CAD inhibition in an *in vivo* setting, we generated previously-activated CD8+ T cells from P14 transgenic mice and pretreated them with PALA before adoptively transferring them into mice which were then infected with LCMV-Armstrong. The percentage of Thy1.1+ cells in the blood of these animals were monitored via cheek bleed on days 4, 6, and 8 post-infection. PALA-pretreatment decreased the speed with which the previously-activated cells expanded in response to antigen encounter (Figure 4D). Thus, in both *in vitro* and *in vivo* settings, inhibition of *de novo* pyrimidine synthesis in previously-activated CD8+ cells impaired their recall response.

As further validation of these findings, we utilized the shRNAs described in Figure S1 to generate CAD-deficient previously-activated CD8+ T cells which were then subject to both *in vitro* and *in vivo* rechallenge. Similar to PALA-pretreatment, introduction of shRNA against the *Cad* gene following T cell activation but before rechallenge impaired cytokine production upon rechallenge (Figure 4E). Adoptive transfer of CAD-deficient antigen-specific CD8+ T cells into mice which were then challenged with infection led to the impairedexpansion of those cells (Figure 4F). Taken together, these findings demonstrate that genetic ablation or pharmacologic inhibition of CAD in previously-activated CD8+ T cells impairs their ability to respond rapidly and robustly to antigen encounter, both *in vitro* and *in vivo*.

### Overexpressing CAD improves CD8+ recall responses *in vitro* and *in vivo*

Thus far, our studies demonstrate that either pharmacologic inhibition of CAD or its genetic deletion leads to decreased recall responses. As such, we next sought to increase CAD activity in previously-activated CD8+ T cells by retroviral overexpression of CAD. We cloned the wild-type *Cad* gene sequence into the GFP-containing MigR1 retroviral backbone to generate CAD-MigR1, allowing us to transfect and visualize cell populations with increased CAD expression. To confirm overexpression, previously-activated CD8+ T cells were generated and transduced with CAD-MigR1 or empty vector, with CAD overexpression increasing both total CAD protein levels and pCAD levels at Ser1859 (Figure 5A). To confirm that this increase in active CAD also led to increased flux into *de novo* pyrimidine synthesis, we again employed amide-labeled ^15^-N glutamine tracing. Previously-activated CAD overexpressing CD8+ T cells demonstrated increased flux into *de novo* pyrimidine synthesis both at rest and upon rechallenge (Figure 5B). When these cells were rechallenged and analyzed for cytokine production *in vitro*, CAD overexpressing cells produced more cytokines and were more polyfunctional when assayed for IFNγ and TNFα production than empty vector controls (Figure 5C).

**Figure 5:**
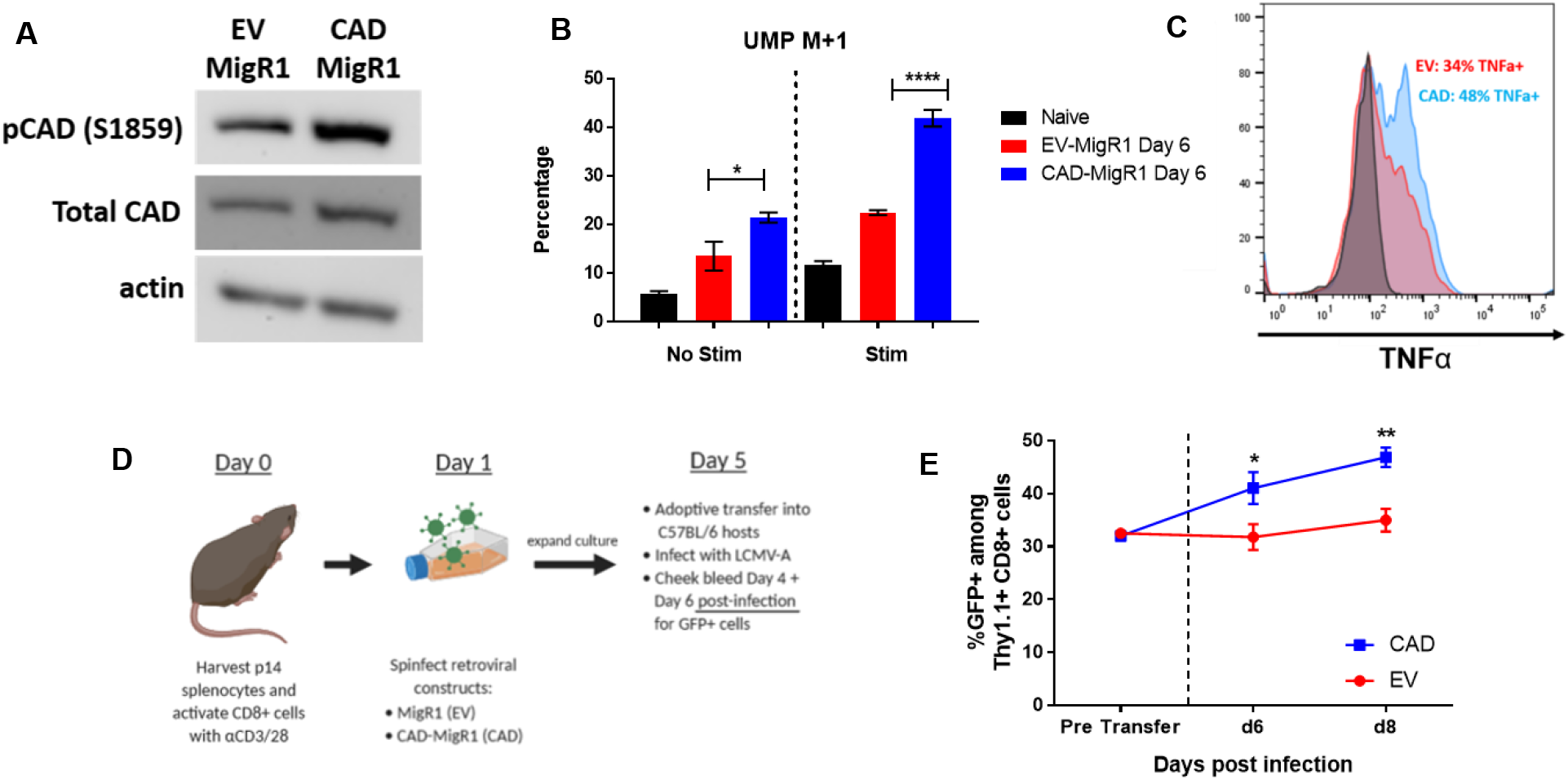
CAD overexpression improves recall response in CD8+ T cells. (A) Western blot analysis of pCAD (S1859) and total CAD levels in CD8+ cells following retroviral transduction with EV-MigR1 or CAD-MigR1 constructs. CD8+ cells isolated from naive C57BL/6 mice were stimulated with α-CD3/28 antibodies for 24 hours before introduction of indicated retroviral constructs with cells analyzed on Day 6 of culture. (B) Percentage M+1 peak data for *de novo* pyrimidine synthesis metabolites measured by LC/MS. EV-MigR1 and CAD-MigR1 cells were generated as in Figure 5A. On Day 6 of culture, freshly-isolated naïve, EV-MigR1 and CAD-MigR1-cultured cells were rested in culture media before transfer to glutamine-free media supplemented with 4mM amide-labeled ^15^-N glutamine, with a portion of each condition stimulated with α-CD3/28 antibodies. C) Cytokine response of retrovirally-transduced cells generated as in Figure 5B following 4 hours stimulation with PMA/ionomycin. Data gated on live CD8+GFP+ cells. Representative flow plot for TNFα positivity is shown, non-stimulated cells in grey. (D) Schematic of adoptive transfer experiments with previously activated P14 retrovirally-transduced cells. (E) Percentage GFP positivity before transfer and at Day 4 and Day 6 following adoptive transfer and infection with LCMV-Armstrong. Data in A are representative of three independent experiments. Data in B are mean ± SD of triplicate samples and representative of two independent experiments. Data in C are representative of three independent experiments. Data in E are representative of three independent experiments with n = 4 to 5 mice per condition. Data in B analyzed with unpaired Student’s t-test. Data in E analyzed with unpaired Student’s t-test at each indicated time point. *p <0.05, **p<0.01, ****p<0.0001.

To explore the effects of CAD overexpression in an *in vivo* setting, we employed CAD-MigR1 or empty vector to generate populations of previously-activated gp33-specific CD8+ T cells with equal percentages of GFP-positivity. These cells were then adoptively transferred into hosts immediately infected with LCMV-Armstrong (Figure 5D). In response to antigenic challenge, the frequency of GFP+ cells in the CAD-overexpression condition increased, while the frequency of GFP+ cells remained largely unchanged in empty vector controls (Figure 5E). This preferential expansion of CAD overexpressing cells provides evidence *in vivo* of a superior recall response when the ability of CD8+ T cells to conduct *de novo* pyrimidine synthesis is enhanced.

### Modulating CAD activity does not alter mRNA transcript levels but affects production of ribosomal RNA precursors

To further elucidate mechanisms for the defect in cytokine production observed in PALA-pretreated cells, we generated previously-activated cells and rechallenged them as before, isolating RNA for qPCR analysis of cytokine transcripts. Intriguingly, CAD inhibition did not impair the magnitude of CD8+ recall response at the level of transcription (Figure 6A). As CAD inhibition consistently impaired cytokine protein production as demonstrated in Figure 4, we therefore reasoned that the defect in pyrimidine synthesis caused a post-transcriptional inhibition of protein synthesis in these cells. Nucleotide availability has been demonstrated to preferentially affect the production of ribosomal RNA over mRNA in cancer cells (*20*), leading us to hypothesize a preferential role for ribosomal RNA synthesis in resting previously-activated and memory CD8+ T cells. Mammalian RNA polymerase I (Pol I) transcribes a 13kb pre-rRNA, a polycistronic precursor later processed into the mature 18S, 28S, and 5.8S ribosomal RNA species (Figure 6B). As the precursor rRNA molecule is relatively short-lived and rapidly undergoes post-transcriptional processing and maturation, the detection of regions undergoing rapid processing such as the 5’ externally transcribed spacer region (5’ETS) can be used to assess active pre-rRNA synthesis (*21,22*). PALA-pretreatment decreased pre-rRNA as determined by qPCR using primers for 5’ETS (Figure 6C). These observations suggest that the role of CAD-induced *de novo* pyrimidine synthesis in previously-activated and memory T cells is not to enhance cytokine transcription, but rather to enhance the synthesis of ribosomal RNA.

**Figure 6:**
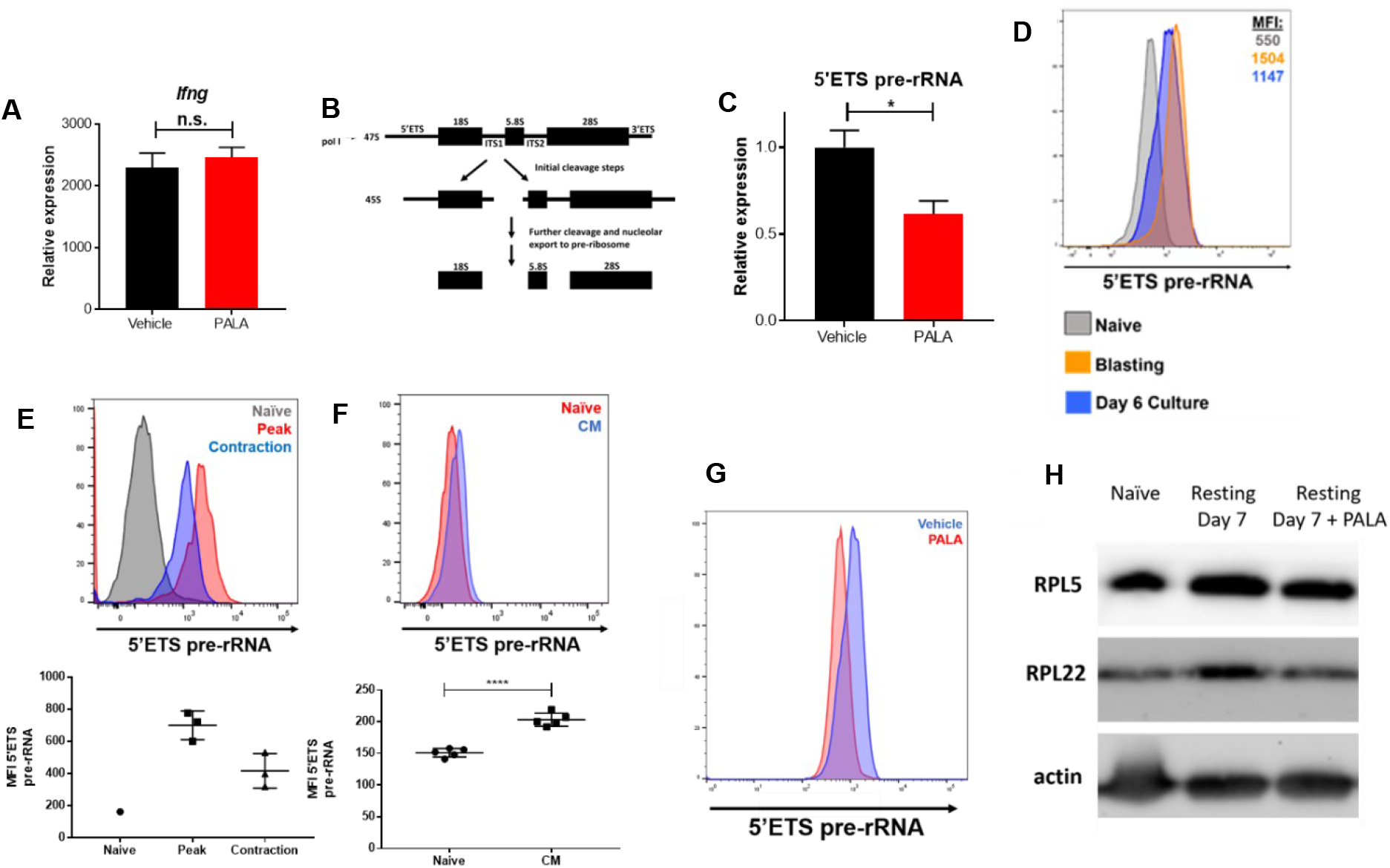
Inhibition of de novo pyrimidine synthesis restricts rRNA synthesis without affecting mRNA production. (A) Relative expression of *Ifng* transcript in previously activated CD8+ cells treated with Vehicle or PALA. Cells were generated as in Figure 4A and stimulated with α-CD3/28 antibodies on Day 6 of culture. Expression relative to resting Vehicle treated cells, β-actin used as housekeeping gene. (B) Diagram of the 45S pre-rRNA molecule transcribed by RNA Polymerase I before processing into 18S, 28S, and 5.8S ribosomal RNA molecules. (C) Relative expression of 5’ETS pre-rRNA in previously activated CD8+ cells treated with Vehicle or PALA. Cells were generated as in Figure 4B, expression relative to Vehicle treated cells, β-actin used as housekeeping gene. (D) 5’ETS pre-rRNA mean fluorescence intensity plotted from naïve, blasting, and resting CD8+ cells. Naïve C57BL/6 CD8+ cells were isolated with a portion immediately fixed for PRIME Flow® analysis. An additional portion was isolated at 48 hours (Blasting) with the remainder expanded in media with IL-7 and IL-15 for an additional four days before isolation (Day 6 Culture). (E) 5’ETS pre-rRNA mean fluorescence intensity in CD8+ cells responding to Vaccinia-OVA infection. Naïve OT-I CD8+ cells were isolated with a portion immediately fixed for PRIME Flow® analysis and the remainder adoptively transferred into C57BL/6 hosts which were then infected with Vaccinia-OVA. Splenocytes were harvested at Day 8 (Peak) and Day 15 (Contraction) and Thy1.1+ CD8+ cells were interrogated for 5’ETS pre-rRNA signal by PRIME® Flow (n=3 mice per condition). (F) 5’ETS pre-rRNA mean fluorescence intensity in naïve and central memory phenotype cells of uninfected mice. C57BL/6 splenocytes were harvested and naïve cells (Live CD8+CD44-CD62L+) and CM cells (Live CD8+CD44+CD62L+) were interrogated for 5’ETS pre-rRNA by PRIME® Flow (n=5). (G) 5’ETS pre-rRNA mean fluorescence intensity plotted from previously activated Vehicle and PALA treated cells. Cells were generated as in Figure 4B before fixation for PRIME Flow® analysis. (H) Western blot analysis of previously-activated CD8+ T cells generated as in Figure 4A and interrogated for indicated ribosomal proteins. A, C, D, G, and H are representative of three independent experiments. E and F are representative of two independent experiments. Data in A and C show mean ± SD of technical duplicates analyzed by unpaired Student’s t-test. Data in F show mean fluorescence ± SD of 5’ETS pre-rRNA in indicated cell populations analyzed by unpaired Student’s t-test. *p <0.05, ****p<0.0001.

Assessment of mRNA and pre-rRNA on a per-cell basis is not possible with PCR-based techniques. The PrimeFlow® RNA Assay, an *in situ* hybridization assay utilizing branched-DNA technology and which is compatible with flow cytometry, has been demonstrated to specifically and robustly amplify signals from mRNA and miRNA species (*23,24*). Expanding the potential of this technology, we designed PrimeFlow® probes to target the 5’ETS of pre-rRNA to quantify the rates of rRNA synthesis in different cell populations on a per-cell basis. Radiolabeling experiments from the 1960s established that naïve lymphocytes synthesize limited amounts of ribosomal RNA at rest but rapidly induce synthesis during the first few days after activation (*16*). Consistent with these findings, PrimeFlow® analysis of pre-rRNA showed low levels of rRNA in naïve CD8+ cells which were elevated in blasting cells at 48 hours post-activation *in vitro* and persisted at six days of resting in culture (Figure 6D). We next exploited this flow cytometry-based assay to determine if we could detect persistence of pre-rRNA synthesis *in vivo*. Indeed, antigen-specific CD8+ cells analyzed during the contraction phase of response to an infection retained elevated pre-rRNA levels compared to naïve cells, although the signal was lower than that observed at the peak of infection (Figure 6E). In a separate set of experiments and in a manner similar to that described in Figure 3A, pre-rRNA was enriched in central memory-phenotype CD8+ cells (CD44^+^ CD62L^+^) over naïve CD8+ cells (CD44^-^ CD62L^+^) in uninfected mice (Figure 6F). These findings suggest a continuation of pre-rRNA synthesis in memory CD8+ cells beyond the initial burst of rRNA synthesis that is known to accompany the activation of naïve T cells.

Lastly, we tested the effects of CAD inhibition on pre-rRNA production and ribosomal biogenesis. PALA pretreatment of previously-activated cells decreased pre-rRNA detected by PrimeFlow® before and after rechallenge relative to vehicle treatment, in alignment with our qPCR data (Figure 6G). PALA pretreatment of previously-activated cells also decreased detectable ribosomal proteins, consistent with the finding that inhibition of pre-rRNA synthesis leads to reduced ribosomal protein content (Figure 6H). Taken together, these findings demonstrate a role for *de novo* pyrimidine synthesis and active CAD for the generation of rRNA and ribosomal biogenesis. These pathways play a role not only at initial activation, but also continuously in previously activated/memory T cells, facilitating the robust recall response upon rechallenge.

### Human memory CD8+ cells are enriched in phosphorylated CAD and modulating CAD activity affects their recall response

Finally, we wanted to determine if the ability of CAD to promote ribosomal biogenesis in memory T cells was also critical to human T cells. To this end, we analyzed PBMCs from human blood. pCAD was enriched in the CD45RA-CR45RO+CCR7+ central memory CD8+ population compared to the CD45RA+CD45RO-naïve CD8+ populations in three healthy donors (Figure 7A). Utilizing stable mammalian expression vectors, we transiently transfected human CD8+ T cells with CAD or empty vector using electroporation as diagrammed in Figure 7B. *Cad* overexpressing human CD8+ T cells demonstrate higher basal levels of pCAD than those expressing empty vector (Figure 7C). Cad overexpressing CD8+ T cells produced more cytokine than empty vector controls upon rechallenge (Figure 7D). This increase in cytokine production was abrogated by pretreatment of these cells after transfection with PALA or a human RNA pol I-specific inhibitor BMH-21. BMH-21 has previously been shown to specifically arrest Pol I transcriptional activity without eliciting DNA damage and with negligible effect on Pol II (*34,35*). As the product of Pol I activity is ultimately the generation of the ribosomes themselves, we sought to analyze the ribosomal protein content of memory cells in comparison to naïve cells. Figure 7E demonstrates an enrichment of ribosomal proteins in memory cells compared to naïve cells. These findings suggest a requirement in human CD8+ T cells for active CAD and that modulation of nucleotide availability or direct Pol I inhibition in previously-activated cells prior to rechallenge can affect recall response.

**Figure 7:**
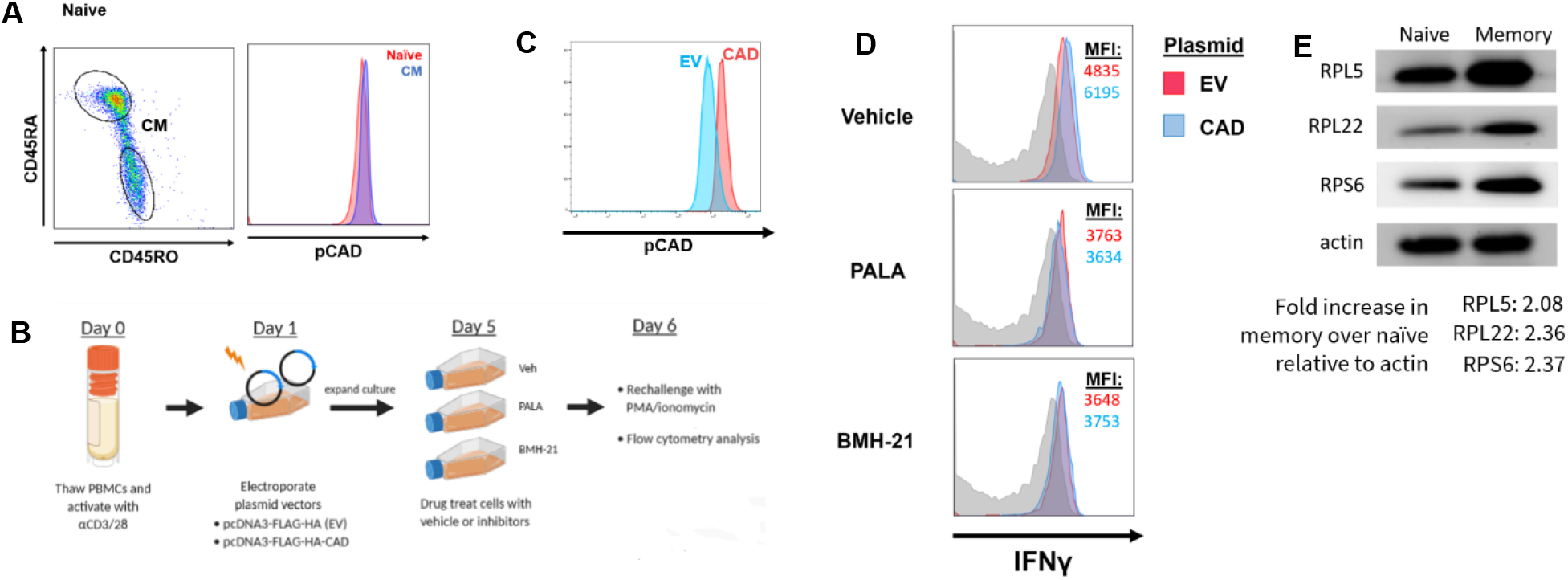
CAD activity in human lymphocytes influences recall response and modulation of pyrimidine synthesis affects ribosomal protein content in human CD8+ T cells. (A) Flow cytometric analysis of pCAD in naïve and central memory human CD8+ T cells. PBMCs were isolated from fresh healthy human blood with an example gating strategy shown, data gated on live CD3+CD8+ cells. (B) Schematic for overexpression of HA-tagged CAD construct in cultured human CD8+ T cells. (C) Flow cytometric analysis of pCAD levels in *Cad*-overexpressing or empty vector-expressing human CD8+ T cells. Data shown are gated on live CD3+CD8+ HA+ cells. (D) Flow cytometric analysis of IFNγ production by CD8+ cells with and without *Cad* overexpression following indicated drug treatment. Cells were exposed to drug for 24 hours before drug removal and four hours of restimulation with PMA/ionomycin. Data shown are gated on CD3+CD8+ HA+ cells. Non-stimulated cells shown in grey. (E) Western blot of ribosomal proteins naïve and memory CD8+ T cells. Human PBMCs were thawed and magnetically sorted into naive (CD8+CD45RO-) and memory (CD8+CD45RO+) populations for lysis and analysis. Data in A representative of three healthy donors. Data in C, D, and E representative of three independent experiments with three frozen PBMC aliquots from three separate healthy donors.

## Discussion

The heightened response of memory CD8+ T cells is one of the hallmarks of adaptive immunity and requires relatively quiescent cells to rapidly reacquire effector function (*1,2,25*). Memory CD8+ T cells emerge from quiescence so quickly that they can divide in as little as two hours after encountering their cognate antigen *in vivo* (*26*). In this study we propose that active CAD in resting memory CD8+ T cells provides a mechanism to support this superior recall response by fueling ribosomal biogenesis. Naïve T cells have been shown to contain low levels of ribosomal proteins and 45S rRNA which increase in an ERK-MAPK-dependent manner following TCR stimulation (*27*). The protein synthetic demands placed on a newly-activated T cell are immense, and it is not unreasonable to expect a concomitant increase in ribosomal biogenesis to provide the necessary protein translation machinery. The role of this process as it pertains to previously-activated cells is not well-studied, although an upregulation of genes involved in ribosomal biogenesis such as those encoding ribosomal subunits and snoRNAs has been reported in memory cells (*28*). The current study both extends the investigation of ribosomal biogenesis beyond the effector phase of a T cell response and provides a mechanism for the superior recall response of memory CD8+ T cells through active CAD and continued pre-rRNA generation.

Although we identify CAD as a downstream phosphorylation target of mTORC1 in CD8+ T cell activation, the maintenance of pCAD in resting cells is mTOR-independent. The precise mechanisms which promote the continued phosphorylation of CAD remain to be determined. Previous studies have identified CAD as a binding partner for 14-3-3ζ (*37*). As members of the 14-3-3 family of proteins are known to sequester targets by binding phosphorylated residues, it is possible that this interaction could protect or maintain a pool of pCAD in CD8+ T cells. Likewise, it is possible that other kinases and inhibition of certain phosphatases may be involved. We can, however, conclude that the mTORC1-dependent reprogramming that takes place during the initial cell activation is integral to the generation of phosphorylated CAD. Along these lines, our findings support the notion that while rapamycin and mTORC1 inhibition can promote memory T cell formation, this is subsequent to the activation mTORC1 during the initial antigen recognition. To this end, our findings are consistent with the role of low dose rapamycin as promoting memory while high dose rapamycin can promote T cell anergy (*13*). To this end, we have previously demonstrated that the specific inhibition of mTORC2 leads to metabolic reprogramming that greatly enhances T cell persistence and hence T cell memory without inhibiting effector function (*6*). Thus, our data strongly support the development of mTORC2 specific inhibitors in order to enhance the efficacy of vaccines. In this way, unbridled mTORC1 activity can promote CAD activation and enhance the speed and robustness of the recall response, while inhibition of mTORC2 leading to enhanced FOXO1 activity and improved memory generation.

Inasmuch as CAD promotes *de novo* pyrimidine synthesis, we initially hypothesized that inhibition of CAD activity would lead to inhibition of cytokine production upon rechallenge at the transcriptional level. To our surprise, inhibition of CAD prior to rechallenge led to decreased cytokine protein without inhibiting mRNA levels. Pursuant to these findings we consequently revealed the role of CAD in promoting rRNA synthesis. Ribosomal biogenesis is among the most energy-intensive cellular processes in eukaryotes, requiring dozens of ribosomal subunit proteins, over 200 different assembly factors and the synthetic activity of all three RNA polymerases to assemble functional ribosomes and export them from the nucleolus (*29*). RNA polymerase I (Pol I), whose sole transcriptional product is the polycistronic pre-rRNA molecule, accounts for over half of all cellular transcription (*30*). The synthesis of ribosomal RNA is the rate-limiting step in ribosomal biogenesis and ribosomal proteins are in fact proteolytically degraded in the absence of appropriate RNA molecules (*36*).

Indeed, our studies suggest that the rapid and robust response of memory T cells is facilitated in part by the continuous generation of ribosomes, such that upon rechallenge, these cells are poised to rapidly respond. As such, our data reveals a novel line of inquiry in terms of studying defective T cell responses, such as T cell exhaustion. Overexpression of CAD and increased flux into *de novo* pyrimidine synthesis facilitated expansion of previously-activated antigen-specific CD8+ T cells in our murine models (Figure 5D) and we propose that a continuation of pyrimidine synthesis is a defining property of memory cells. As cellular therapies with more memory-like transferred cell populations survive longer and control tumors to a greater degree than effector-like populations (*31,32*), increasing CAD activity could have implications in adoptive cell therapy (ACT) or CAR T-cell therapy in clinical settings for treatment of malignancy. Alternatively, inhibition of ribosomal biogenesis could be utilized in the treatment of autoimmune conditions in which the response of memory T cells is known to contribute heavily to pathology, such as multiple sclerosis (*33*). Identification of pre-rRNA synthesis and its relation to ribosomal biogenesis as a potential regulator of T cell memory function therefore invites opportunities to modulate this process in diverse therapeutic settings.

## Materials and Methods

### Study Design

These experiments began as an investigation of CAD phosphorylation downstream of T cell activation. The hypotheses regarding utilization of nucleotides for rRNA came later after qPCR data revealed no difference in cytokine transcript levels under our experimental nucleotide restriction conditions. Unless otherwise stated, laboratory experiments were conducted at least three separate times with representative data shown.

### Mice

Male or female mice between six and nine weeks of age with appropriate age- and sex-matching were used for each experiment. All mouse experiments and procedures were approved by the Johns Hopkins University Institutional Animal Care and Use Committee. C57BL/6, CD4-Cre, OT-I, CD90.1, and P14 transgenic mice were obtained from The Jackson Laboratory. Mice with loxP-flanked *Rptor* and *Rictor* alleles were kindly provided from Mark Magnuson, Vanderbilt University Medical Center, Nashville, Tennessee, USA. *Rptor* -/- and *Rictor* -/- mice with T cells deficient in the indicated protein were backcrossed on to the C57BL/6 background for 9 generations and genotyped according to laboratory guidelines.

### Drugs

Rapamycin was purchased from LC Laboratories. PALA (N-phosphonacetyl-L-aspartate) was provided courtesy of the Developmental Therapeutics Program (DTP) at the National Cancer Institute. BMH-21 was provided courtesy of Marikki Laiho, Johns Hopkins University, Baltimore, Maryland, USA.

### Antibodies and Reagents

The following reagents were purchased or acquired by listed sources or companies:

BD Biosciences: CD8α (53-6.7), CD44 (IM7), IL-2 (JES6-5H4), TNF-α (MP6-XT22), IFN-γ (XMG1.2), CD45RO (UCHL1), CD45RA (HI100), human CD3 (UCHT1), human IFN-γ (B27)

BioLegend: CD62L (MEL-14)

Cell Signaling Technologies: anti-CAD, anti pCAD (S1859, #12662S), anti-S6, anti-pS6 (S240/244, D68F8), anti S6K, anti-pS6K (T389, #9205S), anti-RPL5 (#14568S), anti-RPL22 (anti-β-Actin (D6A8, #8457).

eBioscience: Fixable Viability Dye eFluor 780 (65-0865-14), eBioscience Brand Fixation/Permeabilization set (88-8824-00).

GE Healthcare: Ficoll Paque

Peprotech: Recombinant IL-2, IL-7, and IL-15 cytokines

ThermoFisher: anti-RPL22 (# PA5-97192), PRIME Flow® #CPF-204 Assay ID VP47VTP “5 Prime ETS” bDNA Alexa Fluor 647

Stimulatory anti-CD3 (2C11) and anti-CD28 (37.51) purified in-house from hybridoma supernatants

### Flow Cytometry

Flow cytometry experiments were performed using a FACS Celesta (BD Biosciences) and analyzed using FlowJo software (v.10.3). Intracellular staining for cytokines, pCAD, and pS6 was performed with BD Cytofix/Cytoperm Fixation and Permeabilization solution (BD Biosciences). For PRIME Flow® studies, samples were prepared in accordance with the provided protocol (ThermoFisher).

### qPCR

Total RNA from cells was extracted with TRIzol Reagent (Life Technologies 15596018) RNA was converted to cDNA using the ProtoScript II Reverse Transcriptase (New England BioLabs M0368). qRT-PCR was performed with PowerTrack SYBRGreen Mix (ThermoFisher). 5’ ETS pre-rRNA, Ifng, and β-actin probes were ordered from IDT: 5’ ETS: F 5’-TTTTGGGGAGGTGGAGAGTC-3’ | R 5’-AGAGAACTCCGGAGCACCAC-3’ Ifng: F 5’-CGGCACAGTCATTGAAAGCCTA-3’ | R 5’-GTTGCTGATGGCCTGATTGTC-3’ β-actin: F 5’-CATTGCTGACAGGATGCAGAAGG-3’ | R 5’-TGCTGGAAGGTGGACAGTGAGG-3’ Reactions were run using an Applied Biosystems StepOnePlus 96-well Real-Time PCR. ΔΔCt values were normalized to β-actin and a control group of interest.

### Cell Cultures

T cells were stimulated with soluble anti-CD3 (5 μg/ml) and anti-CD28 (2 ug/ml) for 48 hours, followed by a 10-fold media expansion with IL-2 (1 ng/ml) or IL-7 (10 ng/ml) and IL-15 (20 ng/ml) for an additional 4 days. For MigR1 transfection, cells were spinfected with polybrene and viral constructs at 2500 rpm for 90 minutes at 24 hours into culture. Media was changed 24 hours later to remove polybrene. At the conclusion of culture, live cells were separated by density gradient and then restimulated by incubation with irradiated APCs from C57BL/6 mice that had been pre-loaded with indicated concentrations of SIINFEKL peptide or by PMA (10ng/ml) and ionomycin (1μg/ml) in the presence of GolgiPlug (BD Biosciences) for 4 hours.

### Human T cell experiments

Thawed human PBMCs were plated at 5 million/ml and activated with αCD3/28 antibodies. Twenty-four hours into culture, 1 million cells were suspended in 30ul of EP buffer (Lonza) with 3μg of either pcDNA3-FLAG-HA (EV) or pcDNA3-FLAG-HA-CAD (Manning laboratory) for use in a Lonza Amaxa 4-D Nucleofector. Cells were nucleofected per protocol and replated at 1 million/ml in fresh media containing IL-2 for an additional four days before analysis.

### *Cad* Overexpression Experiments

The CAD-MigR1 retroviral construct was prepared by digesting 2μg MigR1 plasmid (Addgene) with 1ul EcoRI and 1ul XhoI (NEB) at 37 degrees for one hour. HA-FLAG-CAD (Manning laboratory) was digested with amplified with primers to introduce recognition sites for the EcoRI and XhoI cuts made in MigR1. The resulting product was incubated with DH5α, allowed to grow overnight at 37 degrees, then isolated using a Monarch® Plasmid Miniprep Kit (NEB). The isolated products were then sequenced to ensure proper addition of the CAD gene sequence to the MigR1 backbone.

### *Cad* Knockdown Experiments

Naïve T cells were stimulated with soluble anti-CD3 (5ug/ml) and anti-CD28 (2ug/ml) for 48 hours before introduction of 1.6ug empty vector or *Cad*-targeted shRNA (TRCN0000032550, Sigma) using Lipofectamine 3000. Cells were cultured for an additional 24 hours, then washed and plated in fresh media for an additional 3 days of culture before analysis.

### Western Blotting

Cell were lysed at indicated time points in whole cell lysis buffer (RIPA). Blots were developed with SuperSignal West Pico PLUS Chemiluminescent Substrate (Thermo Fisher Scientific, 34578). The images were captured using an UVP BioSpectrum 500 Imaging System and analyzed with the accompanying VisionWorks Pro software.

### Infectious Models

In adoptive transfer (AT) experiments with naïve cells, C57BL/6 CD90.2^+^ host mice were injected intraperitoneally with 10^6^ PFU vaccinia-OVA and administered 5 x 10^5^ naïve CD8^+^CD90.1^+^ (Thy1.1^+^) OT-I T cells by retro-orbital injection. Detection of adoptively transferred OT-I+ cells based on CD8^+^CD90.1^+^ staining. For AT experiments with previously-activated cells, C57BL/6 CD90.2+ host mice were injected intraperitoneally with 10^6^ PFU vaccinia-OVA and administered 5 x 10^5^ CD8^+^ CD90.1^+^ (Thy1.1^+^) OT-I T cells cultured as described in the Cell Cultures section. These cells were expanded with the indicated cytokines and either pre-treated with drug, exposed to shRNA, or transfected with MigR1 retroviral construct.

### Metabolic Flux Tracing

Previously Activated cells were generated by stimulating isolated CD8+ cells from naïve C57BL/6 mice with α-CD3/28 antibodies for 48 hours before expanding media with IL-7 and IL-15, with 1mM PALA added for the final 24 hours of culture in the Previously Activated + PALA condition. Culture contents were then subjected to a Ficoll gradient with additional naïve C57BL/6 CD8+ cells isolated on Day 6 of the experiment. Targeted metabolite analysis was performed with liquid-chromatography tandem mass spectrometry (LC-MS/MS). Metabolites from cells were extracted with 80% (v/v) methanol solution equilibrated at −80 °C, and the metabolite-containing supernatants were dried under nitrogen gas. Dried samples were re-suspended in 50% (v/v) acetonitrile solution and 4ml of each sample were injected and analyzed on a 5500 QTRAP mass spectrometer (AB Sciex) coupled to a Prominence ultra-fast liquid chromatography (UFLC) system (Shimadzu). The instrument was operated in selected reaction monitoring (SRM) with positive and negative ion-switching mode as described. The optimized MS parameters were: ESI voltage was +5,000V in positive ion mode and –4,500V in negative ion mode; dwell time was 3ms per SRM transition and the total cycle time was 1.57 seconds. Hydrophilic interaction chromatography (HILIC) separations were performed on a Shimadzu UFLC system using an amide column (Waters XBridge BEH Amide, 2.1 x 150 mm, 2.5μm). The LC parameters were as follows: column temperature, 40 °C; flow rate, 0.30 ml/min. Solvent A, Water with 0.1% formic acid; Solvent B, Acetonitrile with 0.1% formic acid; A non-linear gradient from 99% B to 45% B in 25 minutes with 5min of post-run time. Peak integration for each targeted metabolite in SRM transition was processed with MultiQuant software (v2.1, AB Sciex). The preprocessed data with integrated peak areas were exported from MultiQuant and re-imported into Metaboanalyst software for further data analysis (e.g. statistical analysis, fold change, principle components analysis, etc.).

## Acknowledgements

M.D.C. designed and performed experiments, analyzed data, and wrote the preceding manuscript. S.S., E.T., I.M.S., J.W., and M.L.A. assisted with acquisition of data. L.Z. performed mass spectrometry experiments. M.M. assisted in the cloning of the CAD retroviral construct. M.L. provided reagents and provided input on experimental design. J.D.P. supervised all studies, contributed to experimental design, and assisted in writing the manuscript. We acknowledge our funding sources including NIH Grant #5R01AI077610-08. We would like to acknowledge Brendan Manning of the Harvard T.H. Chan School of Public Health for providing the wild-type CAD plasmid. We thank Im-Hong Sun and Chirag Patel of the Bloomberg Kimmel Institute for Cancer Immunotherapy at the Johns Hopkins University School of Medicine for consulting about experimental design and review of the manuscript.

## Supplementary Figures

**Figure S1.**
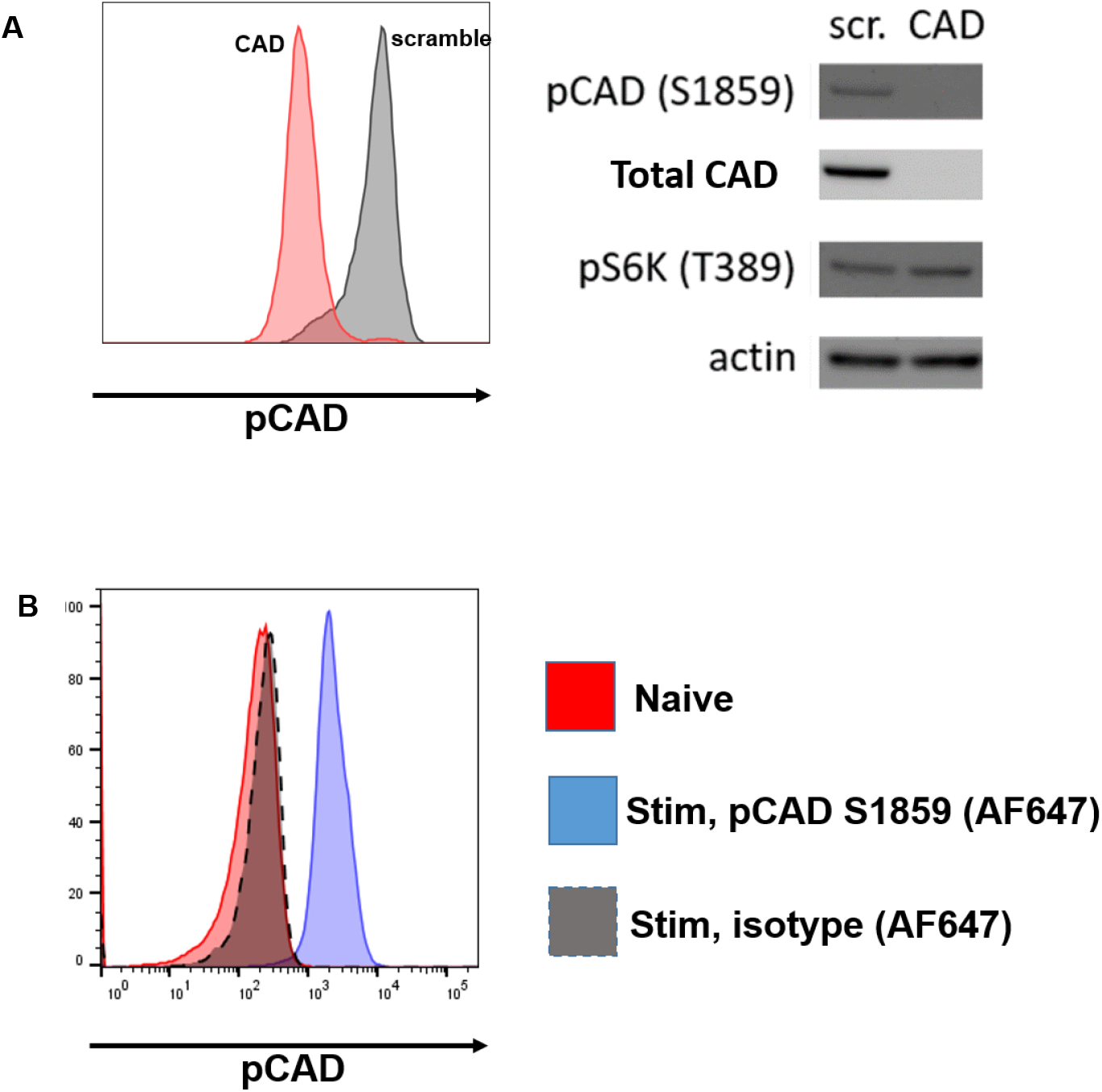
(A) Flow cytometry and Western blot analysis of CD8+ T cells following lentiviral knockdown of *Cad* or treatment with a scrambled control (scr.) CD8+ T cells were isolated from spleens of naïve C57BL/6 mice and stimulated with α-CD3/28 antibodies for 24 hours before addition of indicated lentivirus with polybrene. Cells were incubated for another 24 hours before media was changed to culture media + IL-7/15 for an additional four days of culture before analysis (B) Flow cytometry analysis of C57BL/6 murine CD8+ T cells stimulated for 24 hours with α- CD3/28 antibodies and stained with indicated antibody or isotype control. Data in A and B are representative of three independent experiments.

**Figure S2.**
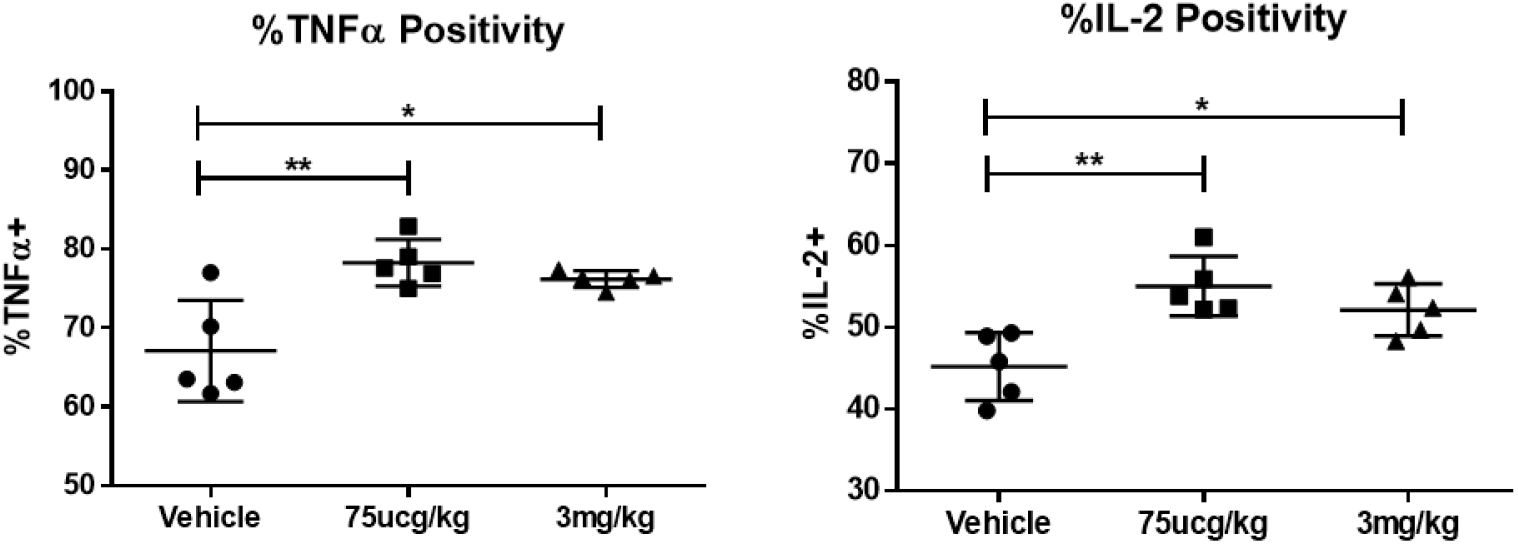
Cytokine positivity from antigen-specific CD8+ cells as described in Figure 3E. Data show mean fluorescence intensity values ± SD analyzed by unpaired Student’s t-test and are representative of three independent experiments. *p <0.05, **p<0.01.

**Figure S3.**
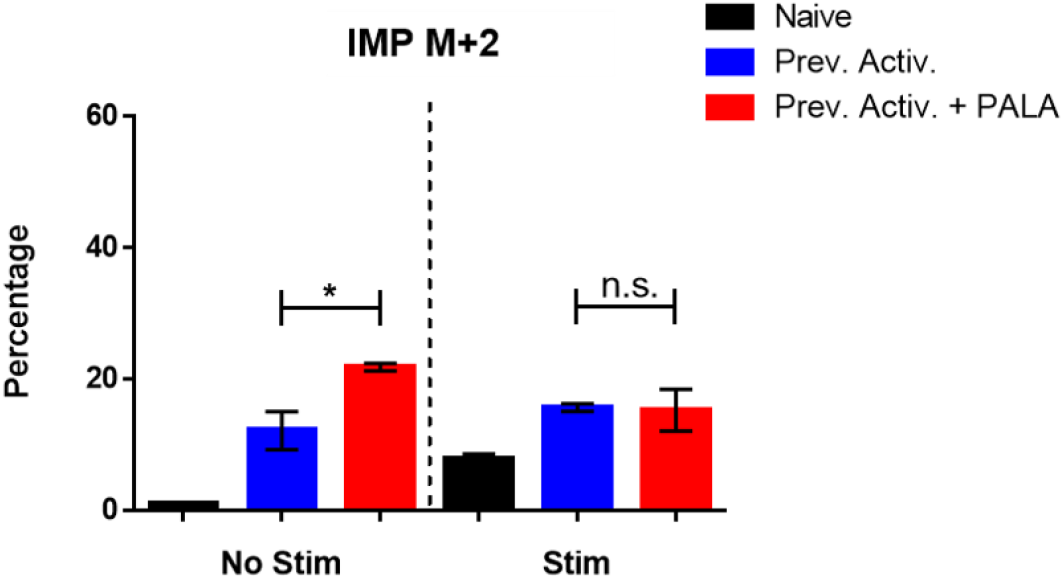
Percentage M+2 peak data for purine synthesis metabolites measured by LC/MS. Cells prepared as described in Figure 2C. Data show mean ± SD analyzed by unpaired Student’s t-test and are representative of triplicate samples analyzed from two independent experiments. *p<0.05, ns = not significant.

